# Directed Nucleosome Sliding in SV40 Minichromosomes During the Formation of the Virus Particle Exposes DNA Sequences Required for Early Transcription

**DOI:** 10.1101/426452

**Authors:** Meera Ajeet Kumar, Karine Kasti, Lata Balakrishnan, Barry Milavetz

## Abstract

Simian Virus 40 (SV40) exists as chromatin throughout its life cycle, and undergoes typical epigenetic regulation mediated by changes in nucleosome location and associated histone modifications. In order to investigate the role of epigenetic regulation during the encapsidation of late stage minichromosomes into virions, we have mapped the location of nucleosomes containing acetylated or methylated lysines in the histone tails of H3 and H4 present in the chromatin from 48-hour post-infection minichromosomes and disrupted virions. In minichromosomes obtained late in infection, nucleosomes were found carrying various histone modifications primarily in the regulatory region with a major nucleosome located within the enhancer and other nucleosomes at the early and late transcriptional start sites. The nucleosome found in the enhancer would be expected to repress early transcription by blocking access to part of the SP1 binding sites and the left side of the enhancer in late stage minichromosomes while also allowing late transcription. In chromatin from virions, the principal nucleosome located in the enhancer was shifted ~ 70 bases in the late direction from what was found in minichromosomes and the level of modified histones was increased throughout the genome. The shifting of the enhancer-associated nucleosome to the late side would effectively serve as a switch to relieve the repression of early transcription found in late minichromosomes while likely also repressing late transcription by blocking access to necessary regulatory sequences. This epigenetic switch appeared to occur during the final stage of virion formation.

## Introduction

Simian Virus 40 (SV40), a member of the Polyomavirus family of DNA viruses, consists of a 5243 base-pair (bp) double-strand closed-circular genome (1). Like many double-strand DNA viruses, SV40 exists as chromatin with the usual assortment of histones and most of the epigenetic marks typical of eukaryotic chromatin when found within an infected cell (2, 3). However, unlike most DNA viruses, SV40 also exists with a typical chromatin structure when found within the virion (3). The organization of SV40 DNA into chromatin throughout its life cycle raises the interesting possibility that epigenetics plays a critical role in the regulation of the various stages of the SV40 life cycle. In support of this hypothesis, we have previously shown that cellular treatments that affect the level of certain histone modifications in the SV40 virus produced during the course of an infection can result in changes in the infectivity of the virus (4). Moreover, we have also shown that the organizations of nucleosomes in SV40 minichromosomes at 30 minutes and 48 hours post-infection differs significantly from the organization of nucleosomes in the chromatin from SV40 virions (5). Since SV40 virion chromatin differs in organization from late stage minichromosomes and also carries information affecting a subsequent infection, the process of encapsidation appears to play an important role in regulating the chromatin structure and resulting biological activity of the virus particle. In order to better understand the nature of the changes in chromatin structure which occur during encapsidation we have analyzed SV40 virions and minichromosomes for the organization of nucleosomes carrying specific histone tail modifications and RNA Polymerase II (RNAPII) and report here evidence that encapsidation leads to a reorganization of chromatin structure within the regulatory region of the virus and relief of the repression of early transcription.

## Materials and Methods

### Cells and Viruses

Working stocks of SV40 virus for infections, virion chromatin, and SV40 minichromosomes were all isolated from BSC-1 cells (CCL-26) freshly obtained from ATCC. The wild-type 776 and mutant cs1085 SV40 viruses used were gifts from Dr. Daniel Nathans. We have previously described in detail the conditions utilized for cell culture (3)

### Infections and Purification of Minichromosomes

We have recently described in detail the procedures that we used for the preparation of minichromosomes (3). In brief, to prepare SV40 minichromosomes, sub-confluent monolayers of BSC-1 cells were infected with 50 PFU per cell of a working stock of SV40 virus originally prepared at low multiplicity of infection. Cells were incubated for 48 hours at which time nuclei were prepared by washing with non-ionic detergent in a low-ionic strength buffer. The nuclei were extracted in a second low-ionic strength buffer and then purified by sedimentation in a glycerol gradient. Aliquots were collected from the top of the gradient and fractions 3-5 which consist of minichromosomes were pooled. In order to separate minichromosomes from encapsidation intermediates we prepared a step gradient consisting of 200 μl of 50% glycerol at the bottom, layered 200 μl of 30% glycerol on the bottom layer, and then layered 600 μl of 10 % glycerol on the two denser layers. The nuclear extracts were layered on top as for a normal centrifugation and following sedimentation the layers were removed as 200 μl aliquots with the 50% and 30% layers kept separate for subsequent ChIP-Seq.

### Purification of virions and chromatin from virions

The procedures for the preparation of chromatin from wild-type and mutant SV40 virions have recently been described in detail (3, 5). Briefly, chromatin was prepared by disruption of low-multiplicity stock virus. The stock virus was concentrated by centrifugation at 50,000 × g for 35 minutes which pellets the virus. The pelleted virus was resuspended in a low-ionic strength Tris-EDTA buffer and digested at least three times with DNAase I at 37 degrees to remove any cellular or viral DNA which might be present on the surface of the virus. At the end of the digestion period an aliquot was removed and analyzed by submerged agarose qgel electrophoresis to determine whether there was any DNA other than SV40 present. Typically, three treatments with DNAase I was sufficient to remove any external DNA. The nuclease-treated virus was then pelleted through 10% glycerol in low-ionic strength buffer at 50,000 XG for 35 minutes to remove any contaminants freed by the nuclease treatment and to again concentrate the virus. The nuclease-digested and concentrated virus was then resuspended in the same Tris-EDTA buffer as above and further treated with a mixture of dithiothreitol and EGTA to disrupt the chemical bonds holding the viral structural proteins together. Following three rounds of disruption at room temperature for thirty minutes each, the virus preparation was again centrifuged on a glycerol gradient as described for minichromosomes and the same fractions pooled as chromatin from disrupted virions.

### Chromatin Immunoprecipitation

A detailed description of the procedures which were used for chromatin immunoprecipitation (ChIP) have recently been published (3). All of the antibodies used were ChIP validated by their respective vendors. Antibodies (and their vendors) included: RNAPII (05-623, Millipore), hyperacetylated H3 (06-599, Millipore), hyperacetylated H4 (06-866, Millipore), H3K4me1 (07-436, Millipore), H3K4me2 (39141, Active Motif), H3K4me3 (04-745, Millipore), H3K9me1 (ab9045, Abcam), H3K9me2 (ab1220 Abcam), H3K9me3 (ab8898, Abcam), and H4K20me1 (39175, Active Motif). ChIPs were performed using Millipore kits according to the supplier’s protocol. Typically, 10 μl of antibody (10 μg) was bound to a mixture of protein A and protein G agarose in dilution buffer for 4 hours. The agarose with bound antibody was then incubated overnight with SV40 chromatin and the bound chromatin purified as previously described according to the protocol present in the kit. The agarose with SV40 chromatin bound to antibody was sonicated to fragment the chromatin and the fragments bound to the agarose separated from the unbound fragments by low-speed centrifugation. The agarose was washed twice with TE buffer from the kit followed by removal of the bound DNA using the buffer supplied in the kit.

### Preparation of sequencing libraries from ChIP samples of SV40 chromatin

The samples obtained from ChIP analyses were first purified using Zymo Research ChIP DNA Clean and Concentrator columns according to the supplied protocol and eluted in 26 μl H_2_O. In order to determine the percentage of SV40 chromatin immunoprecipitated, 1 μl was PCR amplified along with the starting chromatin used in the ChIP. The remaining sample was dried prior to reconstitution with 25 μl of water during the preparation of the libraries. Libraries were prepared using the New England Biolabs Next UltraII kit according to the protocol supplied with the kit with one exception. All reagent volumes were reduced by half for the preparation of libraries with no apparent effect on library quality or quantity. Libraries were purified on Zymo Research ChIP DNA Clean and Concentrator columns according to the supplied protocol and eluted in 10 μl H_2_O. The libraries were then purified by submerged agarose gel electrophoresis using BioRad certified low-melting molecular biology grade agarose and a band corresponding in size to 200-300 base pairs of library DNA was cut out from the agarose gel, purified on Zymo Research Gel DNA Recovery columns using the protocol and reagents in the kit and eluted from the columns in 21 μl water.

### Next Generation Sequencing (NGS)

Prior to sequencing all of the libraries were analyzed for quality and quantity using an Agilent Bioanalyzer. Libraries which did not meet either the quality or quantity requirements were discarded. The libraries were sequenced on an Illumina MiSeq using protocols and reagents from Illumina in the epigenetics core laboratory at the University of North Dakota. For each combination of SV40 chromatin and antibody targeting a histone modification, a minimum of four libraries were prepared from ChIPs of distinct biological samples for sequencing. The actual number of libraries sequenced for each analysis is indicated in the figure legends. Preliminary quality control analysis of FastQ files was performed using FastQC v.0.11.2 (6). The three prime adapters were trimmed from the reads using scythe v0.981 (7). Quality trimming was carried out using sickle v1.33 (8) with a phred score of 30 as the quality threshold; reads with a length less than 45 bp were discarded. To remove contaminating cellular reads, reads aligning to African green monkey *(Chlorocebus sabeus*1.1) or human (hg19) genomes were removed. The remaining unmapped reads were then aligned to the SV40 genome (RefSeq Acc: NC_001669.1), cut at nucleotide (nt) 2666, using Bowtie2 v2.2.4 (9) for wild-type SV40 or in the case of cs1085 the wild-type sequence with the deletion. Duplicate fragments were removed using the Picard Tools (Broad) Mark Duplicates function. Bam files were filtered to contain only fragments between 100-150bp using an awk script. Replicate bam files were merged together using samtools v1.3.1 (10). Bedgraphs normalized to 1x Coverage were generated from filtered, deduplicated reads using Deeptools v2.5.4 (11). Heatmaps were made using the the Z-scores of the normalized coverage, and displayed using IGV v2.3.52 (12).

## Results

### Organization of histone modifications in nucleosomes present in 48 hour wild-type minichromosomes

At 48 hours post-infection SV40 must coordinately regulate DNA replication, early transcription, and differential late transcription to optimize encapsidation and the resulting generation of new infectious virus. This coordinate regulation presumably involves the interplay of positive and negative regulatory factors, critical enzymes such as RNA Polymerase II (RNAPII), and chromatin structure including nucleosome location and associated histone modifications.

Regulation by a combination of nucleosome location and its associated histone modification(s) would be expected to depend upon both the percentage of the regulated chromatin carrying the modification(s) and the location of the histone modification(s) in the chromatin. For example, activation of transcription might be expected to occur on only a subset of the total chromatin present which has been previously committed to transcription with the percentage of chromatin carrying the modification reflecting that subset of chromatin. The specific location of the modification would also be critical since nucleosomes are thought to regulate biological activity either by acting as a barrier to the binding of other factors in the chromatin or by acting as a scaffold to bring other factors to the chromatin.

In previous publications, we have identified histone modifications which were present in relatively large percentages of SV40 minichromosomes (Table 1) (4, 13). As shown in the table, the percentage of SV40 chromatin carrying a particular histone modification varied from approximately 20% to less than 0.01% with many of the modifications present in the range of 1% to 10%. For comparison we also include in this table the percentage of minichromosomes present at 48 hours PI which were bound by RNAPII which was 1%. Assuming that the 1% of minichromosomes bound by RNAPII is an approximation of the fraction of minichromosomes either actively transcribing or activated for transcription, we would expect that a histone modification present in these same minichromosomes would be expected to be present at approximately the same percentage. From the table it is clear that a number of the histone modifications are present in a range suggesting that they might be playing a regulatory role in the minichromosomes associated with RNAPII. The two histone modifications present in the largest percentages, H3K9me1 (22%) and H3K9me3 (12%), are generally thought to be repressive modifications and would appear to be present in a relatively large fraction of the minichromosomes. Consistent with this, we have previously shown that H3K9me1 repressed early transcription at late times in infection (14). Since most of the histone modifications were present at levels suggesting that they may be involved in either positive or negative regulation, we wanted to know whether the modifications present in SV40 minichromosomes late in infection were enriched at specific sites in the SV40 genome that would likely impact the regulation of transcription at this time or be related to the encapsidation process.

**Table 1:** Percentage of SV40 minichromosomes isolated 48 hours post-infection containing specific histone modifications.

| HH3 | HH4 | H3K4me1 | H3K4me2 | H3K4me3 | H3K9me1 | H3K9me2 | H3K9me3 | RNAPII |
| --- | --- | --- | --- | --- | --- | --- | --- | --- |
| 1.3 | 9.9 | 0.01 | 0.4 | 0.08 | 22 | 0.4 | 12 | 1 |

The position of nucleosomes in SV40 minichromosomes isolated at 48 hours post-infection which contained histones modified on their N-terminal tail and for comparison purposes the position of RNAPII were determined using ChIP-Seq. SV40 minichromosomes were prepared and purified as previously described (5).

The minichromosomes were subjected to ChIP analyses using ChIP-grade antibodies that recognized N-terminal tail histone modifications and antibody to RNAPII and fragmentation of the bound chromatin by sonication (15, 16). DNA was isolated from the samples, libraries were prepared, and the libraries sequenced on an Illumina MiSeq.

Sequencing reads were trimmed to remove indexing sequences, any duplicates were removed, and the reads between 100 and 150 base-pairs in size mapped to the SV40 genome. Duplicate reads were removed to minimize the effects of selective PCR amplification during the preparation of each library. SV40 reads between 100 base-pairs and 150 base-pairs in size were used in order to focus on only nucleosome-sized DNA fragments and also to minimize the variation in the size of the gel purified sequencing libraries which we found could vary depending upon the purification conditions. We have previously taken this approach to determine the location of nucleosomes in SV40 chromatin by micrococcal nuclease digestion (5). To generate heatmaps, the individual biological replicates were pooled, normalized to 1X coverage of the SV40 genome, scaled, and displayed.

The number of biological replicates (at least 4) are shown in the figure legends. In the heatmaps the intensity of the color is directly proportional to the number of reads located at a particular site in the genome. If nucleosomes containing a particular histone modification are specifically located within the SV40 genome we would expect to observe bright bands at only a limited number of sites and much weaker or no bands in other areas of the genome. In contrast, if a histone modification was nonspecifically associated with the minichromosome we would expect to observe bands of similar intensity throughout the genome.

The results obtained from SV40 minichromosomes isolated at 48 hours are shown in the combined heatmaps in Figure 1. From the variation in intensity of bands throughout the genome it is apparent that modified histones and RNAPII are only found in specific regions. Not surprisingly, most of the intense bands were found in the well characterized SV40 early and late extended regulatory region (1) which is indicated in the figure. The other region which contains a relatively intense band corresponds to the site of the SV40 miRNA (17) and the SAS transcription unit (18).

**Figure 1.**
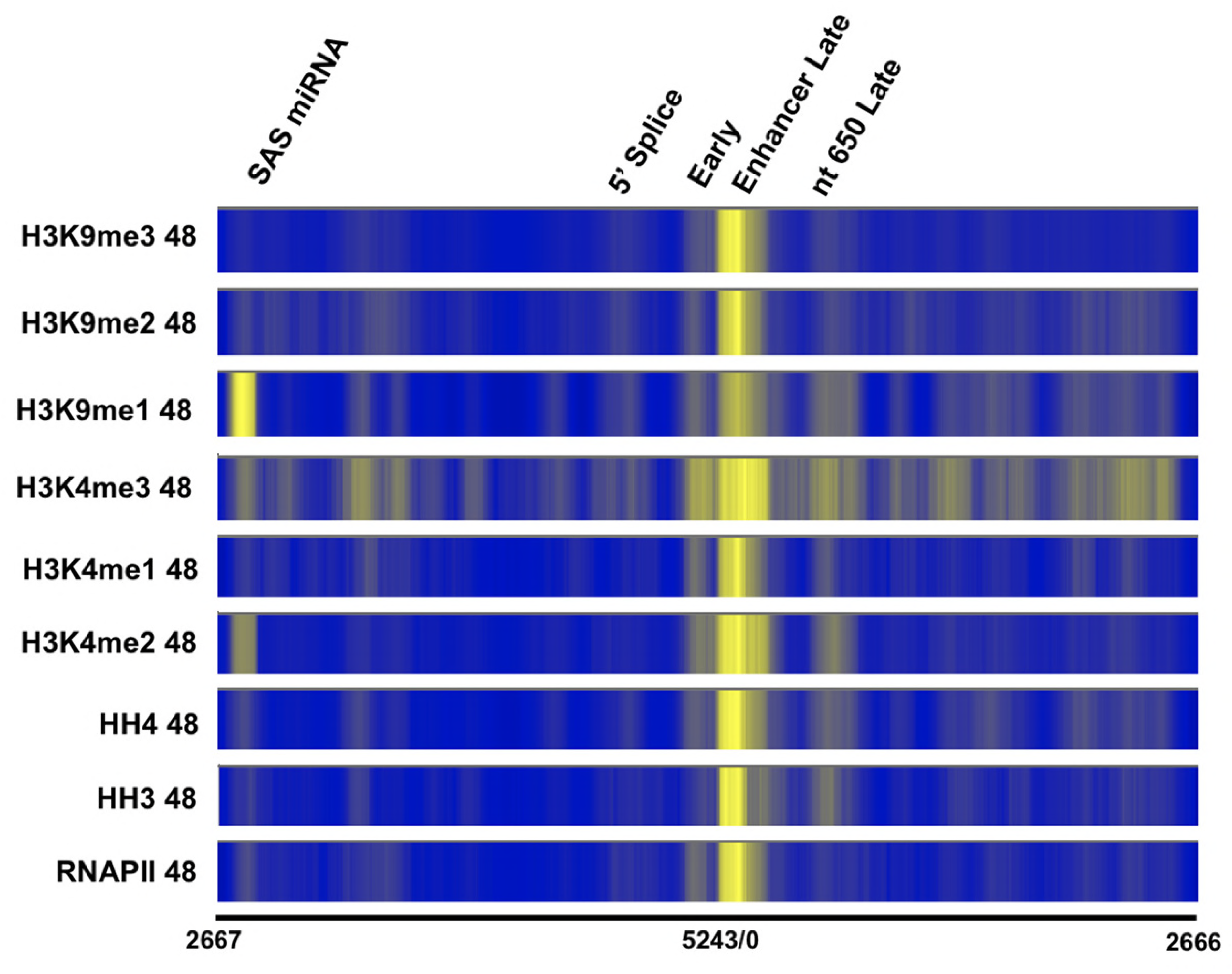
Organization of histone modifications in nucleosomes present in 48 hour wild-type minichromosomes by ChIP-Seq analysis. Minichromosomes obtained 48 h post-infection were subjected to chromatin immunoprecipitation using antibodies to RNAPII and the following histone modifications: HH3, HH4, H3K4me1, H3K4me2, H3K4me3, H3K9me1, H3K9me2, and H3K9me3. The DNA present in the ChIP samples was purified and sequencing libraries prepared using the NEB kit and protocol. The libraries were paired-end sequenced by NGS using an Illumina MiSeq. Following quality assurance and trimming, the reads obtained were mapped to the SV40 wild-type genome which was linearized between nts 2666 and 2667 for display and comparison. The data from at least four biological replicates was merged and heatmaps generated for each ChIP analysis. The actual numbers of biological replecates used were: HH3 (4), HH4 (7), H3K4me1 (5), H3K4me2 (4), H3K4me3 (4), H3K9me1 (8), H3K9me2 (4), H3K9me3 (6), and RNAPII (6). The nucleotide numbers are indicated for orientation with respect to the SV40 genome along with the location of potential regulatory regions.

Within the early and late regulatory region we find the most intense bands for RNAPII to be located within the two copies of the enhancer starting around nt 180 and extending in the late direction to around nt 325 a major late start site. We also observed a less intense band located at the early transcription start site (around nt 5230). The relative intensity of the bands and the absence of bands in the coding regions suggests that the RNAPII is poised for late and early transcription respectively. This would not be surprising since the antibody used to prepare the ChIPs recognizes a peptide containing phosphorylated serine 5 in the CTD of RNAPII which is associated with paused RNAPII.

While the organization of nucleosomes containing the different histone modifications is relatively similar to the results obtained from RNAPII, there are clear individual differences. For example, only HH4, H3K4me1, H3K4me2, H3K4me3, and to a lesser extent H3K9me1 show definite bands over the early transcription start site. Notably HH3 and H3K9me3 appear to be absent at this site. In contrast, all of the histone modifications show relatively intense bands over the enhancer extending into the late start site around nt 325 as seen for RNAPII. Interestingly, while RNAPII does not show a band around nt 650 which is the approximate start of VP1 translation, all of the histone modifications with the exception of H3K9me2 and H3K9me3 show definite bands at this site. Finally, three of the histone modifications, H3K4me1, H3K4me3, and H3K9me1 show relatively intense bands in the region of the SAS and miRNA (indicated on the figure).

In addition to the differences in intensity of the bands for the histone modifications noted above, there also appears to be subtle differences in terms of the specific location of the intense bands. For example, for HH3 and HH4 there appears to be two adjacent sites within the enhancer region of very intense bands. In RNAPII and the other histone modifications it appears that the band on the right is much brighter than the band on the left although the significance of this observation if any is not clear.

Organization of histone modifications in disrupted chromatin from 776 virions In prior studies we have shown that SV40 virions carry biologically-relevant histone modifications (4) and that the location of many nucleosomes in virion chromatin are significantly different than what was observed in 48 hour minichromosomes (5). Together these observations led us to hypothesize that the process of encapsidation to form virions from minichromosomes late in infection might either preferentially utilize certain combinations of nucleosome location and histone modifications in the chromatin as substrates or alternatively function as a chromatin remodeler to prepare the virion chromatin for a subsequent infection. In order to determine the nature of the histone modifications which are retained from minichromosome to virion chromatin and the changes in chromatin structure associated with encapsidation, we have characterized the location of the major histone tail modifications found in chromatin from disrupted SV40 virions.

SV40 chromatin was prepared by disruption of virions as previously described (5), and analyzed by ChIP-Seq using antibodies to HH3, HH4, H3K9me1, H3K9me2, H3K9me3, and H4K20me1. These histone modifications were analyzed because we have previously shown them to be present in a significant percentage of the virion chromatin (4). The results of this analysis displayed as a set of heatmaps are shown in Figure 2. Perhaps not surprisingly, we observed that the most intense band present in virion chromatin for all of the modified histones analyzed except for H4K20me1 appeared to correspond to the most intense band seen in the 48-hour minichromosomes located in the enhancer and late transcriptional start sites between nt 180-325. Similarly, a relatively intense band was located at the early start site for all modifications except HH4 and H3K9me1. In addition, relatively intense bands were also noted for some of the histone modifications around nt 650 which corresponds to the start of the 18.5 S RNA and nt 4885 which corresponds to the 5’ end of the early start site. Finally, there appeared to be a number of other relatively intense bands located at various sites in the SV40 genome for many of the histone modifications which had not been previously seen in the 48-hour minichromosomes.

**Figure 2.**
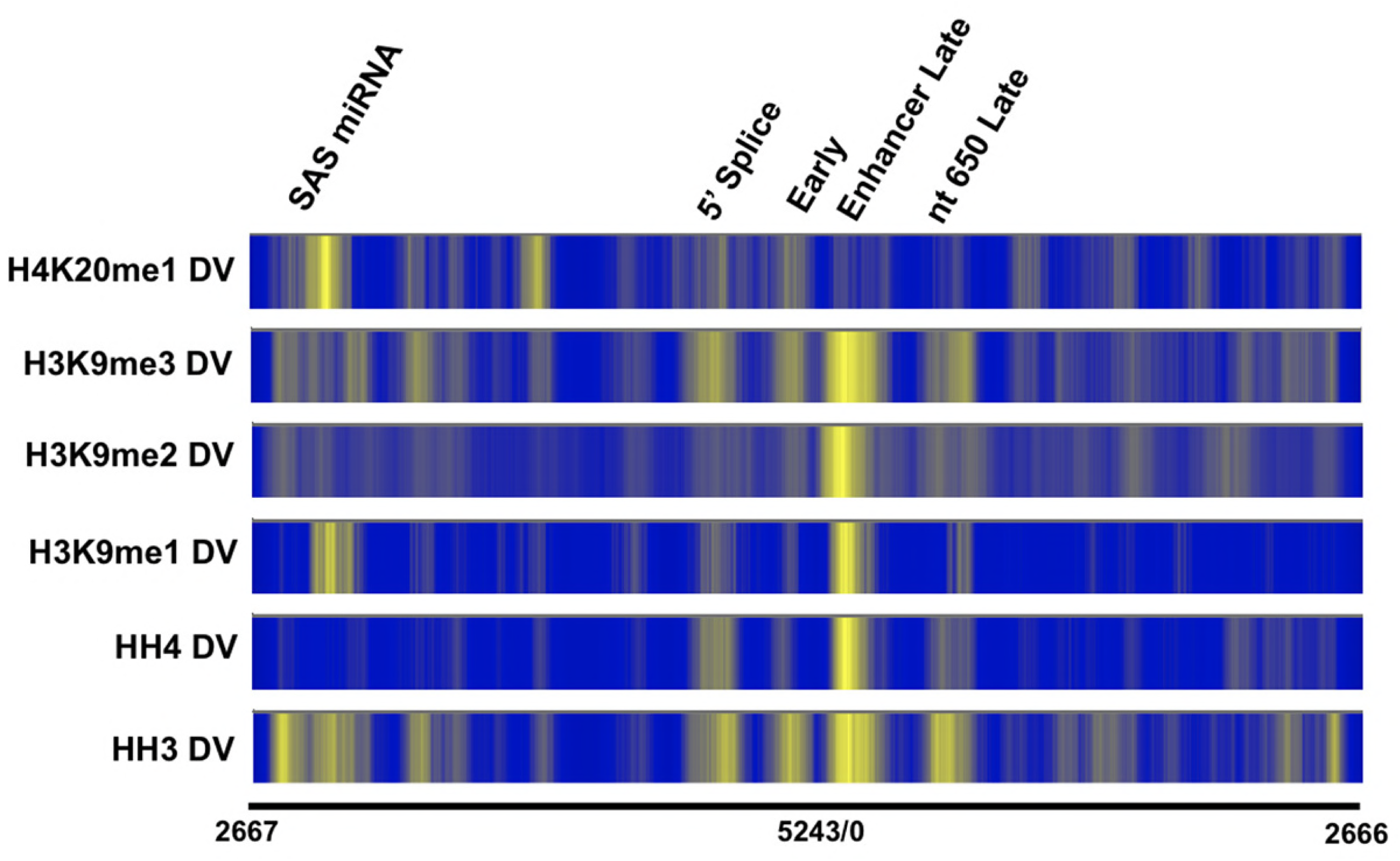
Organization of histone modifications in disrupted chromatin from 776 virions by ChIP-Seq analysis. SV40 chromatin obtained from disrupted virions was subjected to chromatin immunoprecipitation using antibodies to the following histone modifications: HH3, HH4, H3K9me1, H3K9me2, H3K9me3 and H4K20me1. The DNA present in the ChIP samples was purified and sequencing libraries prepared using the NEB kit and protocol. The libraries were paired-end sequenced by NGS using an Illumina MiSeq. Following quality assurance and trimming, the reads obtained were mapped to the SV40 wild-type genome which was linearized between nts 2666 and 2667 for display and comparison. The data from at least four biological replicates was merged and heatmaps generated for each ChIP analysis. The actual numbers of biological replecates used were: HH3 (5), HH4 (5), H3K9me1 (6), H3K9me2 (4), H3K9me3 (5), and H4K20me1 (5). The nucleotide numbers are indicated for orientation with respect to the SV40 genome along with the location of potential regulatory regions.

### Direct comparison of selected histone modifications in 48 hour minichromosomes and virion chromatin

Since we expected that at least some of the 48 hour minichromosomes were serving as substrates for encapsidataion, we hypothesized that for those minichromosomes serving as substrates the banding pattern for their histone modifications should be similar or even identical in the virion chromatin. However, since encapsidation might also drive remodeling we also expected to see some changes such as new bands or shifted bands in the virion chromatin compared to the minichromosomes. In order to determine whether the bands seen in the heatmaps from disrupted virions were located at the same sites as bands present in the 48 hour minichromosome, we directly compared the results for each histone modification from each form of SV40 chromatin as shown in Figure 3.

**Figure 3.**
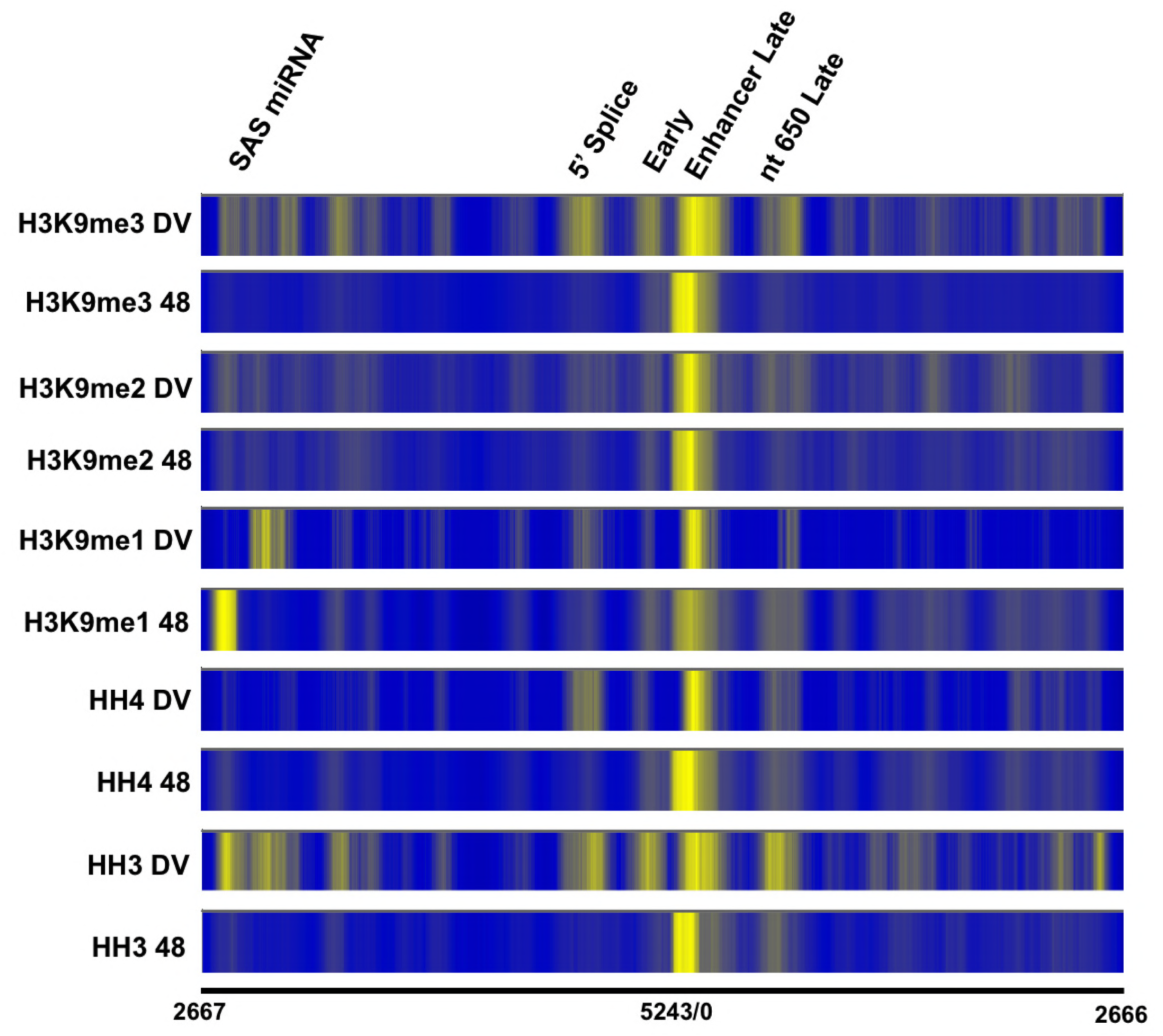
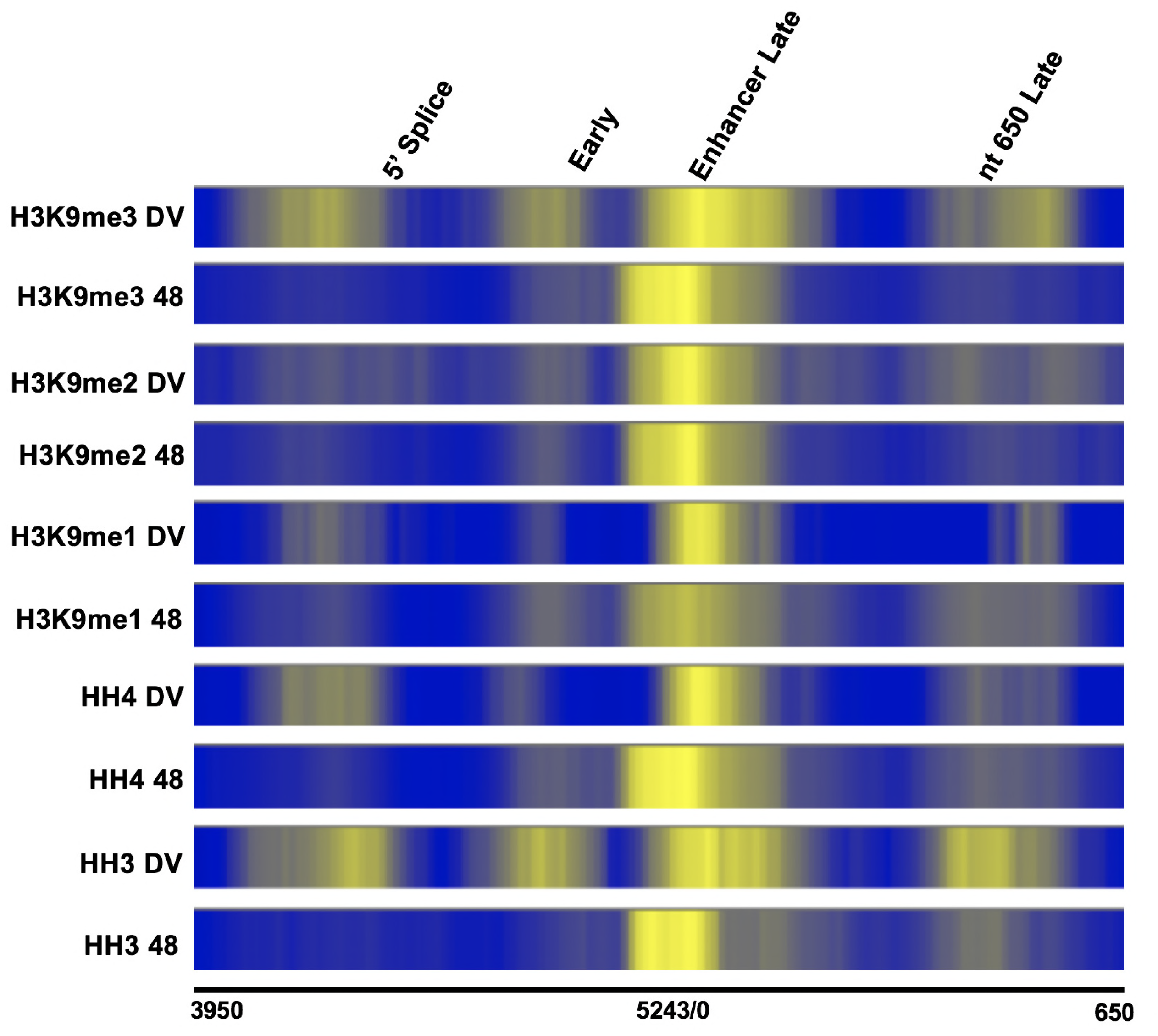
Direct comparison of selected histone modifications in 48 hour post-infection minichromosomes and virion chromatin. The heatmaps of histone modifications present in both the 48 h post-infection minichromosomes and the chromatin from virions from Figure 1 and Figure 2 were similarly scaled and displayed as sets of the same histone modification with the chromatin from disrupted virions above the corresponding chromatin from minichromosomes. The nucleotide numbers are indicated for orientation with respect to the SV40 genome along with the location of potential regulatory regions. In 3A the entire genome is shown while in 3B the regulatory region has been expanded.

The results of this direct comparison between the locations of the major histone modifications present in both forms of SV40 chromatin are shown in the heatmaps in Figure 3. In 3A the entire genome is shown, while in 3B the regulatory region has been expanded to emphasize the differences in nucleosome location. As shown in the figure the major band corresponding to the nucleosome present in the enhancer of the 48-hour minichromosomes was also found in the chromatin from disrupted virions, although with some differences. First, for all of the different forms of histone modification with the possible exception of H3K9me2 which is found in only a small fraction of the chromatin, the major band found in the enhancer of minichromosomes was also present as a major band in the chromatin from virions but appeared to be shifted ~70 bases toward the late side of the genome. This shift in nucleosome position would be expected to change the accessibility of a number of transcription factor binding sites known to affect early and late transcription including SP1 and enhancer binding factors. Second, for most of the histone modifications there appeared to be more bands in the chromatin from disrupted virions than for the chromatin from minichromosomes suggesting a spreading of the histone modifications throughout the genome. This was particularly true for chromatin modified with HH3, HH4, and H3K9me3. Interestingly, the new bands did not appear to be located randomly within the genome but tended to be found at some sites which could potentially be involved in regulation. For example, an increase in band intensity appeared to occur at the early start site, the 5’ early splice site, around nt 650 which was a late start, and around the SAS/miRNA site in chromatin from virions compared to the minichromosomes.

### Heatmap comparison of the location of H3K9me1 in chromatin from disrupted wild-type 776 and mutant cs1085

Based upon our prior studies showing that H3K9me1 plays a critical role in the repression of early transcription by T-antigen (4, 5, 14, 19), we hypothesized that many of the nucleosomes located in the regulatory region which contain H3K9me1 in 48 H minichromosomes and virion chromatin were a consequence of repression of early transcription. In order to confirm that the regulatory nucleosomes containing H3K9me1 were involved in the repression of early transcription, we compared the organization of H3K9me1 in wild-type virion chromatin where repression has occurred to the organization of H3K9me1 in the SV40 mutant cs1085 which lacks T-antigen binding site I and as a consequence does not undergo repression of early transcription (20, 21). We have used this mutant extensively to study the epigenetic regulation of SV40 molecular biology (4, 5, 14, 15, 19, 22, 23) and have shown that the organization of total nucleosomes changes in the regulatory region of the SV40 mutant cs1085 along with a reduction in the proportion of SV40 chromatin containing H3K9me1 (4, 5, 14, 19). It should be noted that in our prior studies (4) although there was a significant reduction in the percentage of SV40 chromatin containing H3K9me1, the percentage never actually went to zero indicating that some nucleosomes containing H3K9me1 remained in the SV40 chromatin.

Comparison heatmaps for the organization of H3K9me1 in chromatin from virions of wild-type and cs1085 are shown in Figure 4. What is clear from the comparison is that the major band corresponding to the regulatory nucleosome on the late side of the promoter has now been shifted significantly in the direction of the late start site. In fact, the nucleosomes corresponding to the brightest bands are likely overlying the major late start site at nt 325. In addition, the relatively weak band located over the early start site in the wild-type chromatin appears to be absent in the cs1085 mutant. These results are consistent with the hypothesis that the regulatory nucleosome located over the enhancer is involved in the process of repression of early transcription by the binding of T-antigen to Site I.

**Figure 4.**
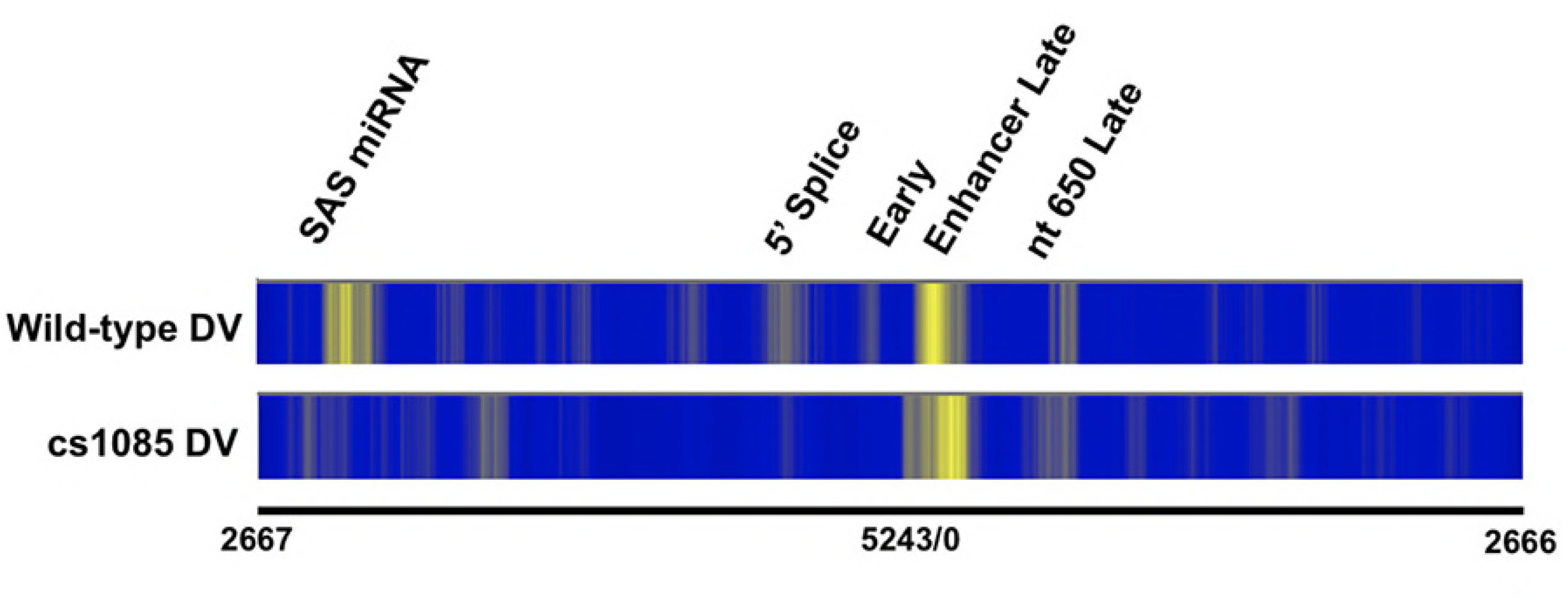
Heatmap comparison of the location of nucleosomes containing H3K9me1 in chromatin from disrupted wild-type 776 and mutant cs1085. Virions from wild-type 776 and cs1085 SV40 were disrupted and the chromatin subjected to ChIP-Seq using antibody which recognizes H3K9me1. Libraries were prepared from the DNA present in the immunoprecipitates and paired-end sequenced using an Illumina MiSeq. The sequence data obtained for the wild-type genome was mapped to wild-type 776. The sequence data obtained for the cs1085 mutant which contains a short deletion was mapped to the sequence of the deletion mutant. For display purposes and comparison to the wild-type results, the sequence data downstream of the deletion was corrected for the size of the deletion by offsetting. For the preparation of the heatmap from the cs1085 disrupted chromatin 4 biological replicates were used. The heatmap for the wild type disrupted virus was the same as in Figure 2. The nucleotide numbers are indicated for orientation with respect to the SV40 genome along with the location of potential regulatory regions.

### Comparison of histone modifications in 776 48-hour minichromosomes and corresponding SV40 encapsidation Intermediates

As shown in Figure 3, there were a number of changes in chromatin structure between 48-hour minichromosomes and the chromatin found in virions. Since the encapsidation process is thought to occur by the accumulation of viral capsid proteins along with compaction of the chromatin, there are three likely ways that the observed changes in chromatin structure could occur. 1, encapsidation could select for a subset of 48-hour minichromosomes that contain the chromatin structure found in the virions. 2, encapsidation could occur on minichromosomes with all or most of the chromatin structure found in the minichromosomes and as encapsidation proceeds the chromatin structure changed. 3, encapsidation could occur on minichromosomes as in 2 but the changes in chromatin structure occur during the final stages of encapsidation after the initial form of the virion, called a previrion, is generated. If mechanism 1 was responsible for the changes we would expect that we would see that the structure found in the virions would also be present in the same relative proportion in the denser encapsidation intermediates. If the changes occurred by potential mechanism 2 we would expect to see a change in proportion between the structure seen for minichromosomes and the chromatin of virions in the encapsidation intermediates. Finally, if mechanism 3 was responsible we would
not expect to see any changes in the encapsidation intermediates since the change in chromatin structure occurred during the final steps of virion formation.

In order to distinguish these three potential mechanisms of chromatin remodeling we analyzed various encapsidation intermediates for the organization of HH3 and H3K9me1. We chose these two modifications because they showed the typical changes in chromatin structure that we had seen for most of the modifications. For this analysis also we took advantage of the previous observation that as SV40 minichromosomes become bound by capsid proteins and begin to condense to form virions they become progressively denser and will sediment further down a glycerol gradient than the minichromosomes. SV40 chromatin was separated on a step gradient consisting of 50% glycerol at the bottom, a 30% glycerol step above, and the rest 10% glycerol. The chromatin in the 50%, 30%, and 10% layers were then subjected to a ChIP-Seq analysis with antibody to either HH3 or H3K9me1 with the results shown as heatmaps in Figure 5.

**Figure 5.**
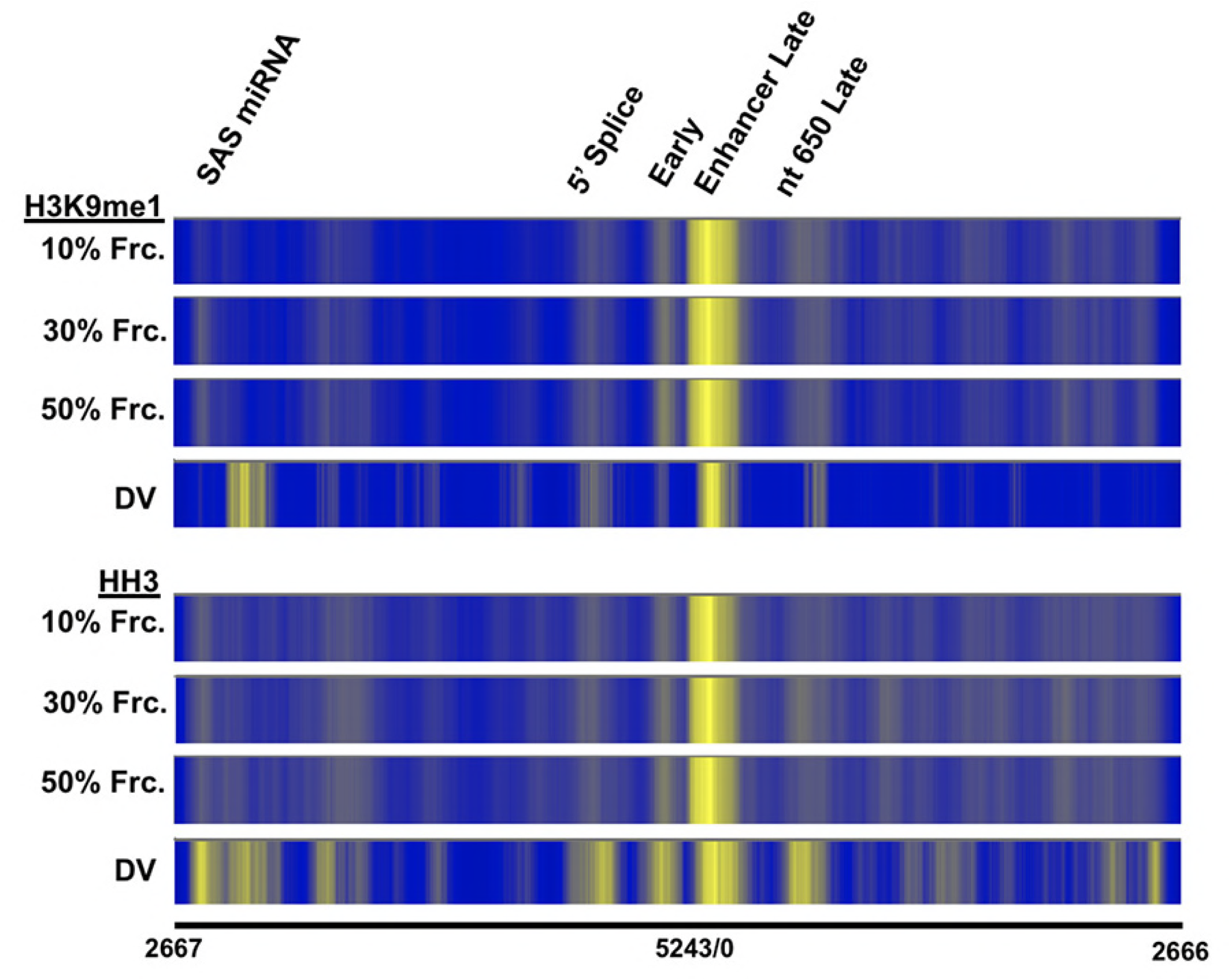
Comparison of histone modifications in 776 48 hour minichromosomes, corresponding SV40 encapsidation Intermediates, and chromatin from virions. SV40 wild-type 776 minichromosomes and encapsidation intermediates were prepared from infected cells by sedimentation on glycerol step gradients. The minichromosomes were found in the 10% glycerol step while encapsidation intermediates of increasing density were found in the 30% and 50% glycerol steps. The minichromosomes and encapsidation intermediates were subjected to ChIP-Seq using antibodies to either HH3 or H3K9me1. Libraries were prepared from the DNA present in the immunoprecipitates and paired-end sequenced using an Illumina MiSeq. The sequence data obtained for each of the samples was mapped to wild-type 776. For each combination of antibody and form of SV40 chromatin four biological replicates were used for the preparation of heatmaps the 10%, 30%, and 50% samples. The heatmap for disrupted virus was the same as in figure 2. The nucleotide numbers are indicated for orientation with respect to the SV40 genome along with the location of potential regulatory regions.

As shown in Figure 5, the patterns of bands for both nucleosomes containing HH3 and H3k9me1 did not change significantly from the gradient fractions containing minichromosomes (10%) to the fractions containing encapsidation intermediates (30% and 50%). The patterns however did differ in the two respects noted above by comparison with chromatin from disrupted virions (DV). Notably the principle band in the regulatory region was not shifted to the position found in the chromatin from virions and the other bands which appear in virion chromatin were still not present in even the densest form of encapsidation intermediate (50%). These results suggest that it is the actual process of forming the capsid of the virion with the SV40 chromatin inside which directs the changes observed between the minichromosomes and chromatin from virions as proposed in hypothesis 3 above.

## Discussion

The transcription starts and by extension the presumed binding sites for RNAPII for the early and late genes (1) and the SAS gene (18) as expected were shown to be associated with RNAPII by our ChIP-Seq analyses in 48 hour SV40 minichromosomes. Since the antibody used to map RNAPII was designed to recognize phosphorylated serine 5 in the RNAPII CTD, it is most likely that the position of RNAPII based upon this antibody reflects the formation of initiation complexes.

Each of the three regions which contained bound RNAPII also contained nucleosomes with specific histone modifications, although importantly the same modifications were not necessarily found at each of the binding sites for RNAPII. At the major RNAPII binding site extending from the enhancer to nt 325, nucleosomes containing more or less all of the tested histone modifications were observed indicating that both positive and negative histone modifications were located in this late transcription regulatory region. Most likely the positive and negative modifications are actually present on different SV40 epigenomes as we previously showed existed at this time in an infection (4) and are related to control of early and late transcription Compared to minichromosomes, the chromatin in virions showed two major changes in structure; 1, certain nucleosomes containing modified histones were shifted slightly including the most intense band in the heatmaps located within the enhancer region and 2, new nucleosomes containing modified histones were present in the chromatin of virions which did not appear to be present in minichromosomes. A schematic representation of the shifting of the enhancer nucleosome is shown in Figure 6.

**Figure 6.**
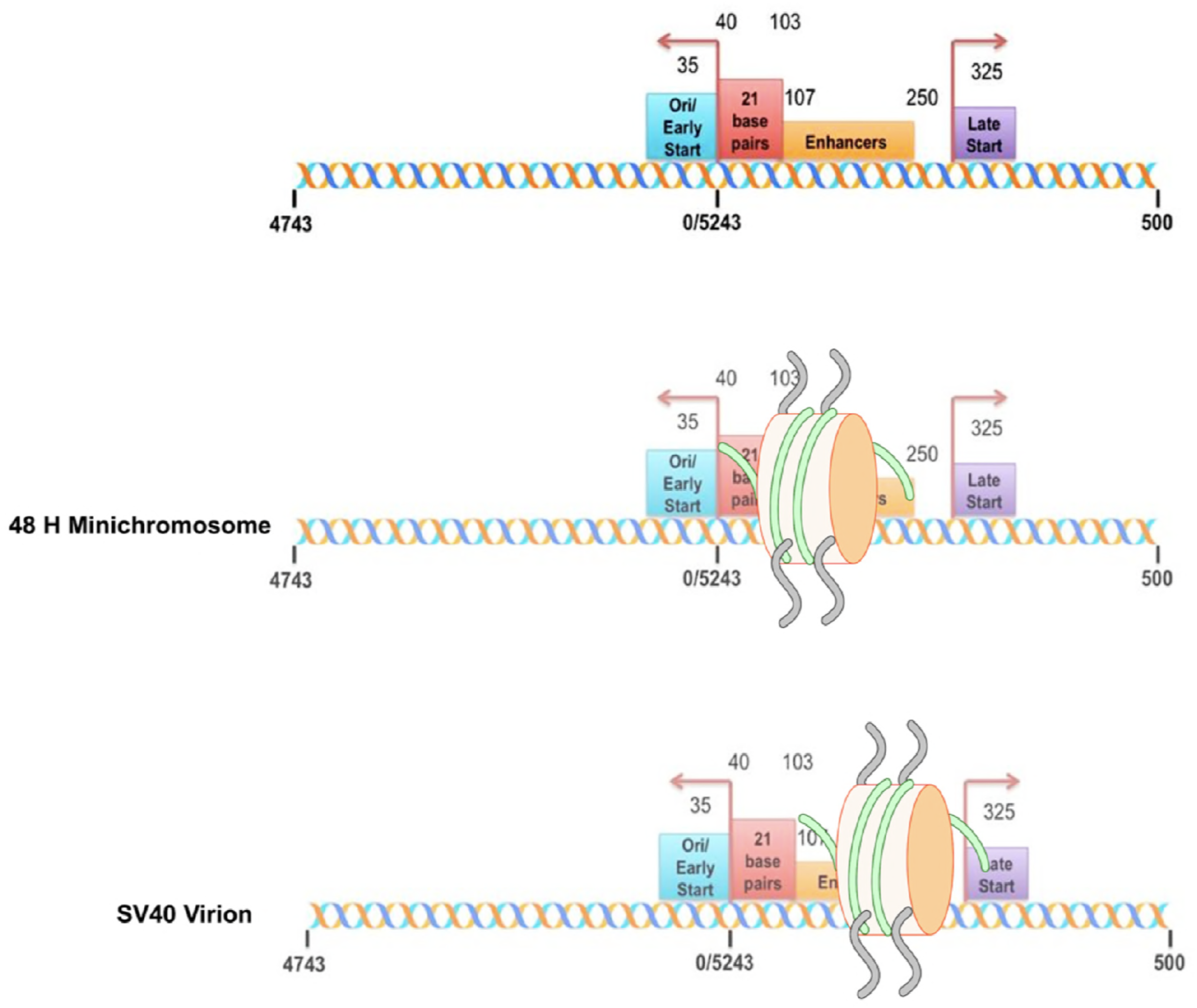
Schematic representation of the position of the regulatory nucleosome carrying various histone modifications found in the SV40 enhancer in 48 hour post-infection minichromosomes and chromatin from disrupted virions. The location of the major regulatory sequences found in the bidirectional SV40 promoter including the 21 bp repeat region, the two copies of the enhancer element, and the early and late transcription start sites are indicated in A. The location of the regulatory nucleosome carrying histone modifications in 48 h post-infection minichromosomes is shown in B and the corresponding location in chromatin from virions is shown in C. The location of the principal regulatory nucleosome was determined using the program R/bioconductor package nucleR v2.20. The center of the nucleosome was extended 72 bp in each direction to determine which regulatory sequences would be protected as shown.

Early transcription requires the TATA, and the 21 bp repeats and is optimized when the intact enhancers are present (24, 25). Since the entire 21 bp repeat region is required for the earliest transcription, the presence of a nucleosome over the GC repeats of III to VI of the 21 bp repeat region as seen in the 48 hour minichromosomes would be sufficient to substantially block this early transcription (25, 26). Consistent with this, we found that this regulatory nucleosome appeared to be positioned at least in part by the repression of early transcription following the binding of T-antigen to binding Site I based upon our result with the mutant cs1085. In contrast, the shifting of this regulatory nucleosome in the SV40 chromatin from the virions would expose the entire 21 bp repeat region along with the 3’end of the first copy of the enhancer and allow for the initiation of early transcription. The minimal late promoter appears to consist of about 68 bases of the enhancer and sequences downstream on the late side (27). The available sequences with a nucleosome positioned as found in the minichromosomes at 48 hours PI may be sufficient to drive late transcription since at least a portion of the critical regulatory sequences would be exposed to binding by regulatory factors (24, 28, 29{Gong, 1988 #518).

We have previously reported on the organization of nucleosomes in SV40 chromatin using micrococcal nuclease digestion (5). We noted the presence of a nucleosome in the enhancer region which was shifted toward the late region in virions although with a reduction in intensity in the digested chromatin.

Presumably, the other changes in chromatin structure which occur during the formation of the virion from late minichromosomes are also related to a shift from a structure that reflects repression of early transcription in the late minichromosomes to one which allows for the initiation of early transcription in the virions. Consistent with this hypothesis we observed the introduction of certain modified histones in nucleosomes at the early transcription start site and at the 5’ end of the early splice site in the virion chromatin.

For the polyomavirus family of which SV40 is a member, encapsidation is thought to occur by a mechanism in which the viral structural proteins associate with the viral chromatin and by crosslinking condenses the chromatin into a virion through encapsidation intermediates of increasing density (30). When encapsidation intermediates were analyzed in our study, we observed that their chromatin was identical to that of minichromosomes and was different from virion chromatin indicating that the formation of the completed virion was responsible for the change in nucleosome positioning. This result supports a model in which those nucleosomes which appear to change position during virion formation do so by a novel encapsidation-associated sliding mechanism.

Although the factors determining nucleosome positioning are relatively well known (reviewed in (31, 32), the locations of specific nucleosomes within genes sometimes change (31). These nucleosomes which occupy close alternative locations are referred to as sliding nucleosomes or fuzzy nucleosomes (31-33). Although fuzzy nucleosomes are not always associated with the direct regulation of transcription as shown here for SV40, a similar type of nucleosome shifting has been described for the human beta-interferon promoter where a nucleosome located over the TATA binding site slides following gene activation (34).

## Acknowledgements

The authors would like to thank the Epigenetic Core Laboratory at the University of North Dakota for sequencing the samples and the bioinformatics analyses. This work was funded by grants from the National Institutes of Health, AI094441 (to B.M.), GM098328 (to L.B) and, GM104360 (to UND Epigenetics Core).

